# Decoding non-conscious thought representations during successful thought suppression

**DOI:** 10.1101/2020.02.02.931352

**Authors:** Roger Koenig-Robert, Joel Pearson

## Abstract

Controlling our own thoughts is central to mental wellbeing and its failure is at the crux of a number of mental disorders. Paradoxically, behavioural evidence shows that thought-suppression often fails. Despite the broad importance of understanding the mechanisms of thought control, little is known about the fate of neural representations of suppressed thoughts. Using functional MRI, we investigated the brain areas involved in controlling visual thoughts and tracked suppressed thought representations using multi-voxel pattern analysis (MVPA). Participants were asked to either visualize a vegetable/fruit or suppress any visual thoughts about those objects. Surprisingly, the content (object identity) of successfully suppressed thoughts was still decodable in visual and executive areas with algorithms trained on perception or imagery. This suggests that pictorial representations of the suppressed thoughts are still present despite individuals reporting they are not. Thought generation was associated with the left hemisphere, whereas thought suppression with right hemisphere engagement. Further, GLM analyses showed that subjective success in thought suppression was correlated with engagement of executive areas, while thought-suppression failure was associated with engagement of visual and memory related areas. These results reveal that the content of suppressed thoughts exist hidden from awareness, seemingly without an individual’s knowledge, providing a compelling reason why thought suppression is so ineffective. These data inform models of unconscious thought production and could be used to develop new treatment approaches to disorders involving maladaptive thoughts.

## Introduction

Selecting and controlling the contents of one’s thoughts is regarded as paramount for achieving goals, learning, controlling emotions as well as psychological wellbeing (Shapiro Jr. and Astin 1998; Østefjells et al. 2017; Carver and Scheier 1982). However, unwanted thoughts can haunt us, leading to outcomes that range from mild discomfort, in the case of thinking about that expensive repair that the house needs, to debilitating mental disorders, such as re-visualizing violent scenes in the battlefield, in the case of post-traumatic stress disorder (PTSD)(Stander, Thomsen, and Highfill-McRoy 2014).

Significant effort has been made in characterizing the behavioural mechanisms of thought control. Seminal work from Wegner and colleagues has shown that, ironically, one’s attempts to suppress thoughts leads to a rebound in the occurrence of the same thought after the suppression effort (Wegner 1994). This, perhaps familiar, outcome has been coined the “ironic process theory”.

Brain imaging has shed light on the areas responsible for thought control. Using a task in which participants had to suppress an object or think freely about anything including the object (Mitchell et al. 2007), the areas responsible for sustained suppression and transient suppression breaks were revealed. Sustained suppression was linked to activations in dorsolateral prefrontal cortex (PFC), while transient activation in bilateral regions of the anterior cingulate cortex (ACC) was observed during the emergence of suppressed thoughts. The ACC has also been shown to be more active during sustained thought suppression compared to free thought (Wyland et al. 2003). In more recent work, suppression and maintenance of visual thoughts was linked to networks encompassing right-lateralized lateral frontolateral areas for suppression (Aso et al. 2016).

In a recent behavioural study, we investigated the sensory traces of suppressed visual thoughts using psychophysics (Kwok et al. 2018). We discovered that the visual representations of suppressed thoughts still lead to perceptual priming, even though the participants reported successful thought suppression. This intriguing result suggests that suppressed visual thoughts might still exist in visual brain areas despite the judgements of successful suppression. In other words, non-conscious visual representations of the thought content might exist in visual cortex, without the individual ever knowing. While previous work has studied the areas responsible for thought suppression, the goal of the present study was to investigate the possible existence of non-conscious thought representations during successful thought suppression.

To investigate the fate of suppressed visual representations, we used a paradigm comparing imagery to suppression. In each trial, participants were instructed to either imagine or suppress the visual thought of a fruit or a vegetable (red apple or green broccoli). This design was adapted from a previous behavioural study in which we used conveniently green or red coloured vegetables and fruits as primes for red/green binocular rivalling gratings (Kwok et al. 2018). Participants reported visual thought intrusions, which we called suppression breaks, when the to-be-suppressed objects appeared in their minds.

As a first goal, we sought to identify the areas engaged by suppression and imagery as well as the areas engaged by successful suppression and suppression breaks. To identify these areas, we used mass-univariate general linear model (GLM) contrasts between conditions as well as multi-voxel pattern analysis (MVPA) to discriminate between tasks in visual regions-of-interests from V1 to V4. Our second aim was to investigate whether successfully suppressed visual and imagined representations could still be decoded despite participants reporting subjective absence of these visual thoughts. To test this, we used MVPA in a cross-decoding generalization design to decode the content of the suppressed thought (apple or broccoli). The generalization design allowed us to reveal whether and where there was representational overlap between perceptual images and successfully suppressed objects as well as between imagined objects and successfully suppressed ones.

Our results show that suppression recruits a right lateralized network, including the prefrontal cortex, the anterior cingulate as well as the superior parietal and temporal cortices. Failure at suppressing visual thoughts (suppression breaks) was associated with enhanced recruitment of memory and higher-level visual areas, while successful suppression was correlated with middle pre-frontal and insula activations. These results are consistent with previous studies in thought suppression using different paradigms thus highlighting the reproducibility and generalization of these effects. Finally, our decoding analysis revealed that, even though participants judged the suppression to be successful, visual perception and visual imagery-like representations were present in the dorsolateral prefrontal and lateral occipital cortex, respectively. These results provide new neural evidence of the pervasiveness of suppressed thoughts and unveil a network of brain areas to be targeted to treat intrusive thought disorders.

## Results

### Imagery/suppression task

Participants were instructed to either imagine or suppresses visual thoughts of an apple or broccoli (Figure 1A, see Materials and Methods for details). Importantly, participants subjectively rated the strength of the visual thought for both imagined and suppressed trials with suppression breaks (Supplemental Figure S1). Suppression vividness ratings (0.85 ±0.17 mean ±SEM) were largely variable among participants, with some participants having very little suppression breaks (e.g., participants 4, 6, 9, 14, Figure 1B), while others reported a large amount of suppression breaks (e.g., participants 1, 3, 12). This was in contrast with the imagery vividness ratings (2.78 ±0.09 mean ±SEM) that were more homogeneous across participants (Figure 1C). This is consistent with the individual differences in thought control revealed in our previous study (Kwok et al. 2018). For analyses exploring suppression success and/or failure (Figures 3 and 4), only participants having at least 25% of suppression breaks or successful suppression were considered, which corresponded to 8 participants (marked with a ★).

**Figure 1.**
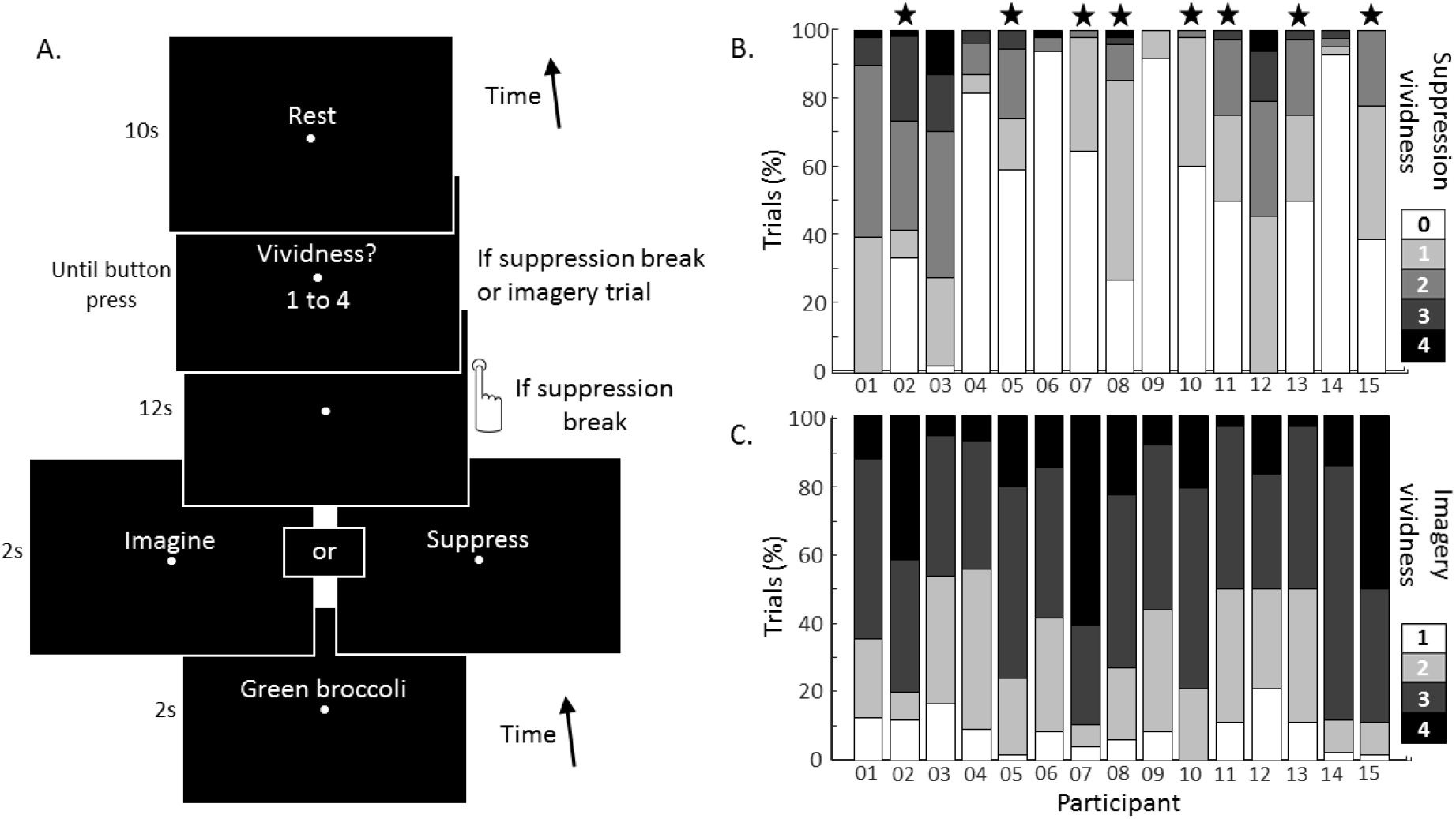
Imagery/suppression fMRI task. **A**. Imagery / Visual thought suppression. Every trial started by a written cue indicating the object to be imagined or suppressed (either a green broccoli or a red apple) for a duration of 2 seconds. After this, the task instruction was presented: “Imagine” or “Suppress”, for 2 seconds. The fixation point remained on the screen for 12 seconds during which the participants tried to either visualize the cued object as vividly as possible or to suppress the visual thought of it. In suppression trials, participants pressed a designated button (same button irrespective of the object to be suppressed) to report a suppression break event, i.e., when the mental image of the object to be suppressed appeared in their minds. In imagery trials and suppression trials with suppression breaks, participants were asked to report the subjective intensity of the visual thought experienced in a vividness scale from 1 (low) to 4 (high). In suppression trials with no suppression breaks, the vividness prompt was not shown and vividness for that trial was assigned to 0. After every trial a inter trial interval of 10 seconds was observed, a fixation point and the word “rest” were displayed on the screen. **B**. Vividness rating in suppression trials for each participant. Suppression vividness from 0 (suppression success) to 4 (highly vivid suppression break) as the percentage of trials for every participant. Participants had a wide range of suppression break ratios. For analyses comparing suppression success and failure, only participants having at least 25% of suppression breaks or successful suppression were considered, which corresponded to 8 participants (marked with a ★). **C**. Vividness rating in imagery trials for each participant. Unlike the vividness ratings for suppression trials, vividness ratings in the imagery trials were more homogeneous across participants. This suggests that the differences across subjects in vividness ratings in the suppression conditions correspond to inter-individual differences in thought-control (Kwok et al. 2018) rather than inconsistences in the vividness report.

### Imagery and suppression engage differently lateralized networks

A GLM analysis (see Materials and Methods for details) revealed a left-lateralized network was associated with imagery production and a right-lateralized network associated with suppression (Figure 2A). This is consistent with previous neuroimaging and lesion studies (Farah 1984; D’Esposito et al. 1997; Aso et al. 2016; Garavan, Ross, and Stein 1999).

**Figure 2.**
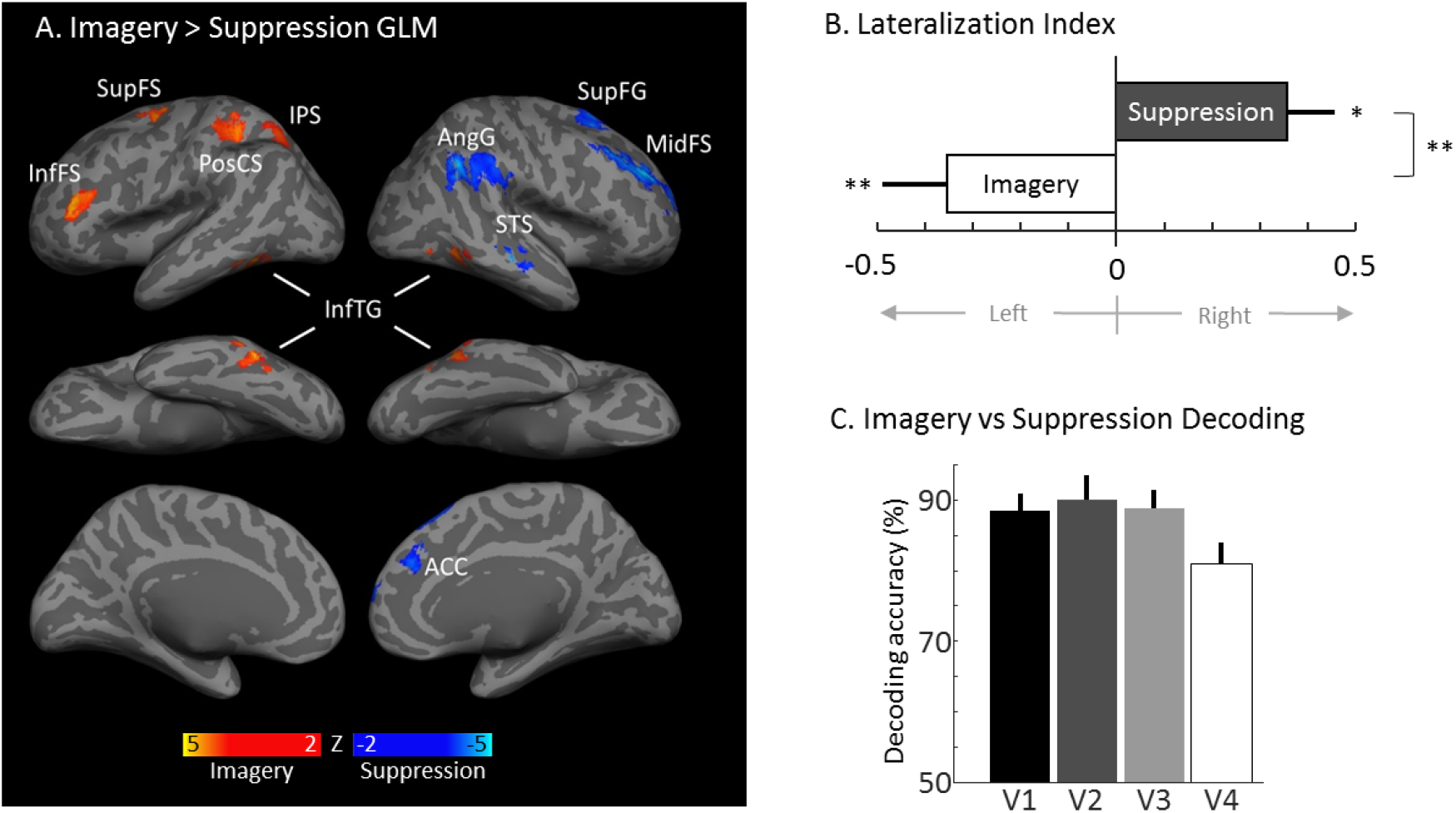
Imagery and suppression engage two different networks. **A**. Imagery > Suppression contrast. Imagery activations (blue) were found in high hierarchy visual areas (InFTG) bilaterally. Left lateralised imagery driven activations were also found on executive areas (InfFS and SupFS) and attention related areas (IPS). Suppression (red), on the other hand, was associated with left lateralized activations in executive (AAC, SupFG, MidFS), high visual (STS) and language (AngG) areas. All activations are p<0.001 (voxel level) and p<0.05 cluster level correction (GRFT) for multiple comparisons. AAC: anterior cingulate cortex; AngG: angular gyrus; InfFS: inferior frontal sulcus; InfTG: inferior temporal gyrus; IPS: intraparietal sulcus; MidFS: middle frontal sulcus; PosCS: post central sulcus; SupFG: superior frontal gyrus; SupFS: superior frontal sulcus. STS: superior temporal sulcus. **B**. Lateralization index for the Imagery > Suppression contrast. Lateralization index as the absolute value of the significant activations across hemispheres (see Methods for details). Imagery activations were predominantly left lateralized, mean −0.35, two-tailed t-test, t(14)=3.57, p=0.003, ci=[0.14, 0.57], uncorrected; consistent with previous reports. Suppression related activations were on the other hand predominantly right lateralized, mean = 0.35 two-tailed t-test, t(14)=−2.61, p=0.02, ci=[−0.64,−0.06], uncorrected. Lateralization indexes for imagery and suppression were significantly different, two-tailed t-test, t(14)=4.1, p=0.001, ci=[0.34,1.08]. **C**. Task decoding. Visual regions-of-interest contained useful information to reliably classifying (above 80% accuracy) imagery from suppression trials, thus indicating that these conditions engage visual areas differently. V1: 88.36%, one-tailed t-test t(13)=14.67 p=10^−6^, ci=[83.73, inf]; V2: 90.04%, t(13)=11.38 p=10^−6^, ci=[83.8, inf]; V3: 88.66%, t(13)=13.59 p=10^−6^, ci=[83.62, inf]; V4: 80.95%, t(13)= 9.7, p=10^−6^, ci=[75.16, inf], all p-values FDR corrected, q = 0.05. Error bars correspond to +1 SEM.

**Figure 3.**
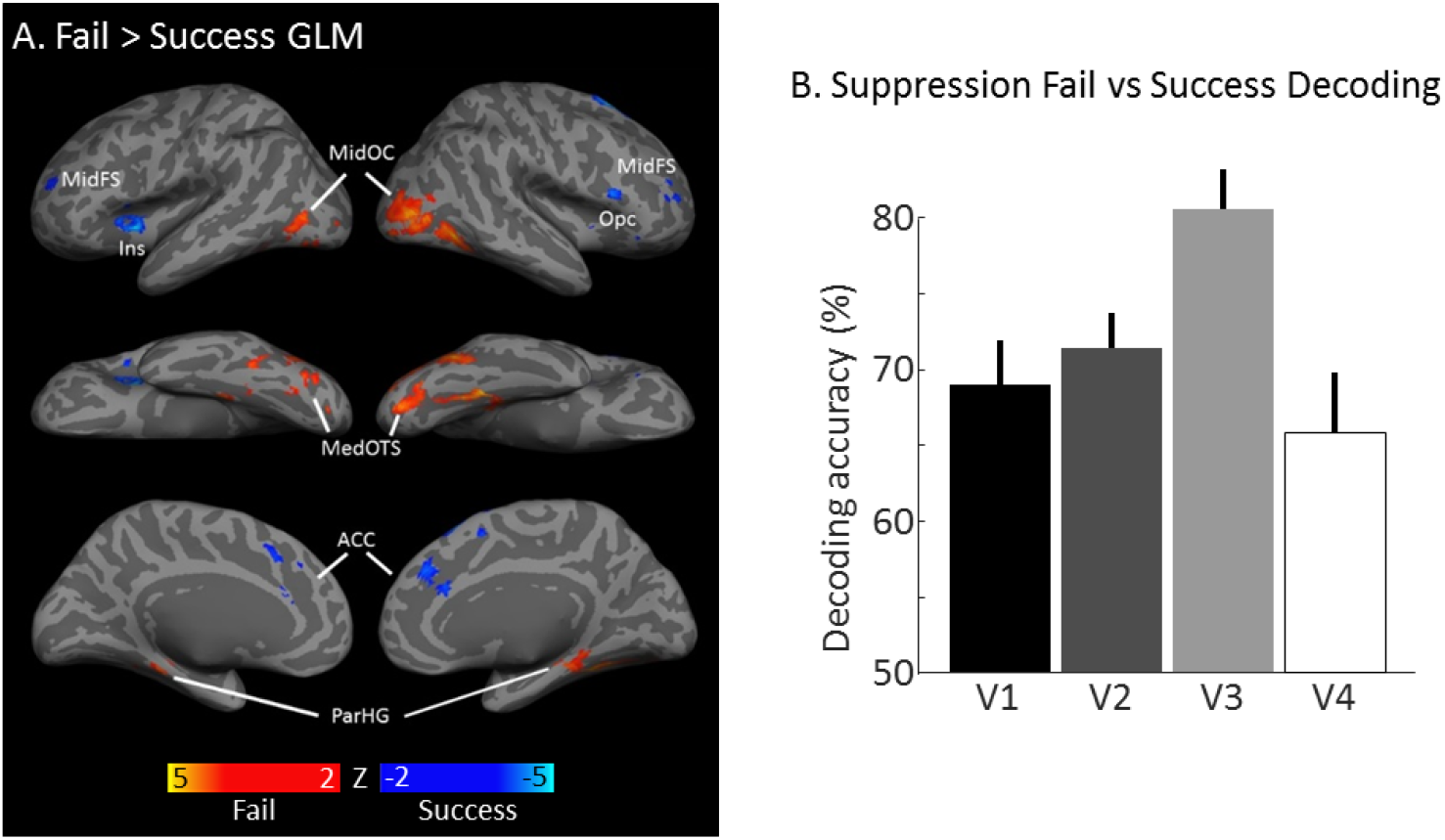
Failed suppression is correlated with activations in visual and memory areas. **A**. Failed > Successful suppression contrast. Failed suppression was associated with posterior activations along the visual stream in areas such as the MidOC, MedOTS and in memory related areas such as the ParHC and the hippocampus (not shown). On the other hand, successful suppression was associated with anterior activations in executive areas such as the MidFS, the Opc and the ACC. This results indicate that control over suppressed thoughts obeys an engagement of executive control areas while failure at suppressing thoughts is accompanied by a hyperactivity of visual and memory related areas. All results are p<0.001 (voxel level) and p<0.05 cluster level correction (GRFT) for multiple comparisons. ACC: anterior cingulate cortex; Ins: insula; MidFS: middle frontal sulcus; MidOC: middle occipital cortex; MedOTS: medial occipito-temporal sulcus; Opc: operculum; ParHC: parahippocampal gyrus. **B**. Task decoding. Visual regions-of-interest contained useful information to classifying failed from successful suppression trials. V1: 68.95% one-tailed t-test t(7)=6.34, p = 2.66·10^−4^, ci=[63.28, inf]; V2: 71.38%, t(7)=9.08, p = 4.13·10^−5^, ci=[66.91, inf]; V3: 80.5%, t(7)=11.3, p= 2.06·10^−5^, ci=[75.39, inf]; V4: 65.87%, t(7)=4.02, p = 0.003, ci=[58.3886, inf], all p-values FDR corrected, q = 0.05. Error bars correspond to +1 SEM.

**Figure 4.**
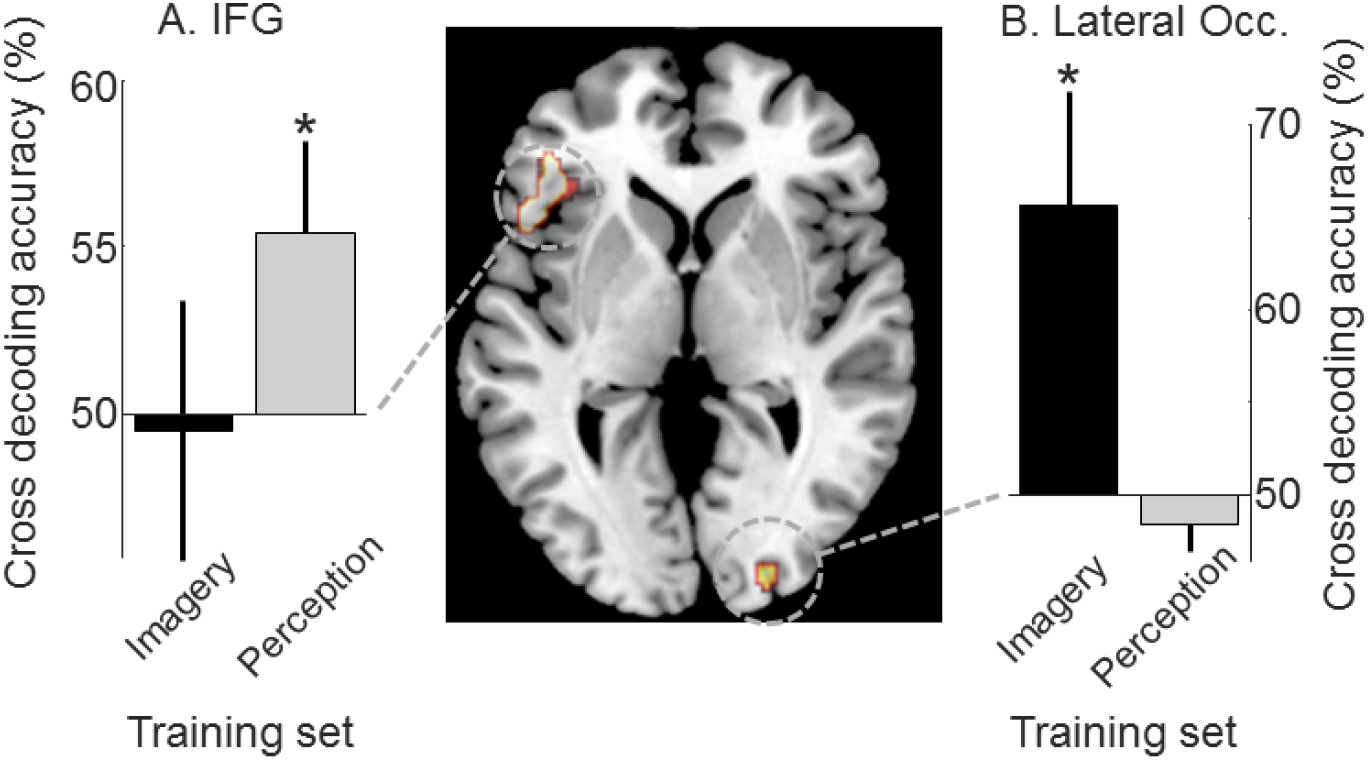
The contents of subjective successful suppression are decodable using information from perception and imagery. To test whether subjectively successfully suppressed thoughts shared informational content with imagery and perceptual representations, we performed a cross decoding analysis. We thus attempted to decode the content of the successfully suppressed thoughts (broccoli or apple) using classifiers trained on imagery trials (black bars) or classifiers trained on perception (grey bars), on regions-of-interest were the contents of suppression were most readily extractable (ROI threshold Z > 2; see Methods for details). The contents of successfully suppressed thoughts were decoded above chance and using patterns from perception trials on the inferior frontal gyrus (**A**. IFG) 55.4% one-tailed t-test t(7)=1.98, p=0.044, ci = [53.9,inf] and using patterns from imagery trials in the lateral occipital (**B**. LOC) 65.6% one-tailed t-test t(7)=2.53, p=0.0198, ci = [50.2,inf]. These results indicate that subjectively successfully suppressed thoughts contain similar information to imagery representations (arguably visual in nature as contained in visual areas) and to perceptual representations in executive areas.

The brain areas activated by both imagery (on the left hemisphere) and suppression (on the right hemisphere) were the superior frontal, prefrontal and temporal cortex (Figure 2A). This is consistent with results highlighting the role of the right prefrontal cortex in inhibitory control (Garavan et al. 2002; see Aron, Robbins, and Poldrack 2014 for a review). On the other hand, the left prefrontal and frontal cortex has been associated with the production of imagery content in different sensory modalities (Lundstrom et al. 2003; Yoo et al. 2003; Sabatinelli et al. 2006).

We found brain areas specifically associated with imagery in the inferior temporal cortex bilaterally (Figure 2A). These areas are known to code for high-level objects representations (Rust and Dicarlo 2010; Grill-Spector 2003) thus, visualization of real-life objects (i.e., apple and broccoli) are expected to engage these areas. On the other hand, suppression specifically engaged right superior temporal areas, specialized in verbal processing. Additionally, suppression was associated with activations in the right anterior cingulate cortex (ACC) an area involved in inhibitory control.

A formal analysis of lateralization (Figure 2B, see Materials and Methods for details) of imagery and suppression across hemispheres revealed that imagery activations were significantly left lateralized (LI=−0.3539, p=0.0031, one-sample, two-tailed t-test against 0, t(14)=−3.5701, ci=[0.1423, 0.5706], uncorrected). On the other hand, suppression driven activations were significantly right lateralized (LI=0.3565, p=0.0204, one-sample, two-tailed t-test against 0, t(14)=2.6146,, ci=[−0.6441,−0.0636], uncorrected).

### Imagery and suppression can be discriminated using spatial patterns of activations in visual areas

While no significant differences in activations in lower-level visual areas (V1 to V4) were detected using GLM univariate analysis (Figure 2A), the informational content in these areas revealed by multivoxel-pattern analysis (MVPA) allowed us to discriminate imagery trials form suppression ones with great accuracy (decoding accuracy > 80%, p = 10^−6^, one sample t-test against 50%, FDR corrected, Figure 2C). MVPA takes into account the spatial pattern of activation associated with imagery and suppression to discriminate between these two tasks rather than relying on overall differences in BOLD response. Therefore, this analysis is more sensitive and better suited to reveal subtle differences that are beyond the sensitivity of univariate analyses (Kriegeskorte, Goebel, and Bandettini 2006). This result shows that the spatial patterns of activations in visual areas differ consistently between imagery and suppression, even though their average activation level does not differ significantly.

Differences between imagery and failed suppression were found in memory, executive and higher level visual areas (Supplementary Figure S2). Imagery related activations were only found in the right hemisphere (inferior temporal and parahippocampal cortices). On the other hand, suppression fail showed noticeably less lateralization compared to the imagery vs suppression contrast. Suppression fail activations were found in both hemispheres (inferior parietal, and frontal areas) except for the right anterior cingulate.

### Failed and successful suppression are associated with posterior vs anterior activations

We employed a GLM analysis to compare activations in failed vs successful suppression trials. This analysis revealed that successful suppression was associated with bilateral activations in the medial prefrontal cortex, the anterior cingulate and the insula (Figure 3A). The engagement of these set of areas is consistent with their involvement in control and monitoring. The medial prefrontal cortex has been implicated in inhibitory control (Aron, Robbins, and Poldrack 2014), whereas the insula is implicated in conflict monitoring (Carter et al. 1998; Gehring and Knight 2000; see Botvinick, Cohen, and Carter 2004 for a review).

On the other hand, failed suppression was associated with activations in visual areas: the middle occipital cortex and the medial temporal cortex. Interestingly, we also found bilateral activations in memory related areas such as the parahippocampal cortex and the hippocampus. The hyperactivity in visual and memory areas during suppression breaks is consistent with the access and generation of visual sensation (Dijkstra, Bosch, and van Gerven 2017; Ishai, Ungerleider, and Haxby 2000) as suppression breaks can be conceptualized as a case of involuntary imagery (Pearson and Westbrook 2015; Pearson 2019).

Using a MPVA analysis, we further found that lower-level visual areas, from V1 to V4 contain information about the success or failure in thought suppression (Figure 3B). Classifiers decoded moderately well the spatial activation patterns associated with failed and successful suppression (p < 0.005, one-sample t-test against 50%, FDR corrected, q = 0.05) with the highest accuracy found in area V3 (decoding accuracy 80.5%). This result shows that reliable information about the success and failure of suppression can be found in lower visual areas.

### Decoding the contents of successful suppression using perceptual and imagery information

In order to investigate the fate of suppressed thoughts, we used a cross-decoding generalization approach to track successfully suppressed representations. To do so, we trained classifiers to discriminate between the perceptual content (green broccoli or red apple) in an independent visual perception task with diverted attention (see Materials and Methods for details), as well as in imagery trials from the main experiment. We then tested SVM classifiers on successful suppression trials (back and grey bars represent imagery-successful suppression and perception-successful suppression cross-decoding respectively, see Materials and Methods for details). Using this analysis, significant above chance decoding accuracy would show that perceptual or imagery representations are generated even though participants rate their suppression as successful.

We found significant decoding of successfully suppressed thoughts content in two brain areas: the left inferior frontal gyrus and the right lateral occipital cortex. The left inferior frontal gyrus (IFG) was found to encode perception-like representations during successful suppression (grey bar, Figure 4A), showing significantly above chance decoding accuracy (55.4% one-tailed one-sample t-test against chance: 50%, t(7)=1.98, p=0.044, ci = [53.9,inf], uncorrected). Evidence of imagery-like representations were however not found in this area (decoding accuracy p>0.05, black bar, Figure 4A). This result suggests that perception-like representations of the suppressed stimuli are still present in executive areas despite the reports of successful suppression.

On the other hand, the right lateral occipital cortex was associated with imagery-like cross-decoding (black bar, Figure 4B). The lateral occipital showed significant decoding accuracy (65.6% one-tailed t-test t(7)=2.53, p=0.0198, ci = [50.2,inf], uncorrected). We however did not find evidence for perception-like representations in this area (decoding accuracy p>0.05, grey bar, Figure 4B). This result suggests that imagery-like representations of the suppressed stimuli in visual areas are still present despite the subjective success at suppressing them.

We further sought to investigate whether perception and imagery patterns would also generalize the representational content of successful suppression in visual ROI (from V1 to V4). We, however did not find significant above chance decoding in functionally defined visual areas (Supplementary Figure S3).

## Discussion

### Suppressed thoughts are still there

Using MVPA, we detected visual perception-like neural representations in the inferior pre-frontal cortex and imagery-like neural representations in the lateral occipital cortex despite subjective reports that the thoughts were suppressed from awareness. A number of previous results have shown that suppressed thoughts influence behaviour despite subjective success at censoring them. Classic studies (Wegner et al. 1987; Wegner 1994) have shown that suppressing thoughts leads to a rebound of thought intrusions after withdrawing suppression efforts, pointing to the pervasiveness of suppressed thoughts. In a recent experiment, we objectively measured the strength of suppressed visual thoughts as a bias on subsequent bistable perception (Kwok et al. 2018). Our results revealed that the perceptual trace of successfully suppressed thoughts was as effective as voluntary imagery at biasing subsequent binocular rivalry. The current results add to these behavioural findings by showing that neural representations of suppressed visual thoughts are still present in the visual cortex and executive areas. This finding sheds light into how successfully suppressed thoughts can interact with perception and affect behaviour, despite thoughts feeling like they have been successfully suppressed out of mind.

### The nature of the suppressed-thought representations

Suppressed thought representations were found to be similar to perceptual and imagery representations. Information patterns associated with visual perception were able to decode the contents of successful suppression, however, the locus of the information was found in the left inferior prefrontal cortex, an area not classically regarded as visual (see however: Hagler and Sereno 2006; Silver and Kastner 2009; Friedman and Goldman-Rakic 1994 for evidence of visual topographic maps in frontal and prefrontal cortices). This result could thus be interpreted as evidence that visual-like representational content of suppressed thoughts in frontal areas might be somewhat abstract, akin to categorical visual representations (Freedman et al. 2001). On the other hand, imagery-like representations were found in the right lateral occipital cortex which is known to house visual object representations (Grill-Spector, Kourtzi, and Kanwisher 2001; Pourtois et al. 2009). Interestingly, imagery representations have been shown to generalize nicely with perceptual representations in the lateral occipital (Cichy, Heinzle, and Haynes 2012), thus suggesting that the information we found in the LOC might also be perceptual in nature. Why suppressed representations did not generalize to perception in the LOC? (Figure 4B, grey bar). One explanation is that even though there could have been representational overlap between perception and suppressed thoughts, classifiers were unsuccessful in cross-decoding due to the difference in signal strength and thus pattern reliability across both modalities (Sterzer, Haynes, and Rees 2008).

Interestingly, classifiers using information contained in retinotopically organized visual areas from V1 to V4 were able to discriminate with great accuracy between imagery vs suppression and between successful and failed suppression. These results indicate that lower visual areas house reliable information about the cognitive state even though we did not find differences in the global level of activation as shown in the GLM analyses.

### Awareness of the suppressed thoughts

While participants reported not having a visual sensation associated with the suppressed objects in the successfully suppressed trials, we cannot however rule out the notion that participants may have failed to report visual intrusions of suppressed thoughts. Interestingly, our previous behavioural study, strongly suggests that participants report intrusions faithfully, as shown by a series of control experiments in a similar suppression/imagery paradigm (Kwok et al. 2019).

Importantly, subjective criteria about the existence or absence of thought intrusions is of clinical relevance to ascertain symptoms associated with psychopathology as well as treatment success (Reynolds and Brewin 1998; Holmes and Mathews 2010; Ehlers and Clark 2000). We thus believe that subjective reports about thought suppression are a legitimate way to measure them and represent a suitable way to relate the present results with clinical research.

### A network of areas as potential targets for therapy of intrusive thoughts

The working assumption in our paradigm is that voluntary visual imagery and visual thought suppression are related processes (Pearson and Westbrook 2015; Pearson 2019; Kwok et al. 2019). While voluntary visual imagery involves the maintenance of a visual objects in one’s awareness, visual thought suppression strives to prevent the selected object from entering awareness. Importantly, failure of thought suppression shares a common phenomenology with voluntary imagery, as conscious visual content is present in both cases. However, brain activations differ between imagery and suppression failure as shown in Supplementary Figure S2.

By comparing imagery to suppression, we were able to identify areas involved in voluntary imagery generation vs visual thought suppression. This analysis revealed two lateralized networks: a left-lateralized network associated with imagery and a right-lateralized network associated with suppression, consistent with previous results (Farah 1984; D’Esposito et al. 1997; Aso et al. 2016; Garavan, Ross, and Stein 1999). These networks included prefrontal, frontal and parietal areas. A body of literature has linked activations in the right prefrontal cortex with several forms inhibitory control, such as error detection, correction and inhibition implementation (Garavan et al. 2002; Garavan, Ross, and Stein 1999; Aron et al. 2003; see Aron, Robbins, and Poldrack 2014 for a review). In contrast, activations in the left prefrontal and frontal cortices have been associated with the production of imagined content across different modalities (Lundstrom et al. 2003; Yoo et al. 2003; Sabatinelli et al. 2006). Suppression also selectively recruited the right ACC, which has been implicated in conflict monitoring (Carter et al. 1998; Gehring and Knight 2000; see Botvinick, Cohen, and Carter 2004 for a review), which engagement would be important to monitor and detect thoughts intrusions during thought suppression.

Imagery selectively recruited the inferior temporal gyrus, an area associated with high-level visual representations (DiCarlo and Cox 2007; Grill-Spector, Kourtzi, and Kanwisher 2001; Grill-Spector 2003), which it is likely to be important to visualize the stimuli.

The failed vs successful suppression contrast highlighted brain areas responsible for adequate control of thoughts vs those associated with thought intrusions. Again, our analyses found the PFC and ACC as areas implicated in thought control or successful thought suppression. This result suggests that engagement of this network of areas is important for maintaining censored suppressed thoughts. On the other hand, thought intrusions were associated with hyperactivity of memory areas: the parahippocampal cortex and the hippocampus as well as visual areas as the lateral occipital and the inferior temporal cortex. This result suggests that during thought suppression a control/inhibition network in frontal areas would oppose the activation of a sensory/memory network located posteriorly. Imbalances in the activation of these networks might have a tangible effect in thought control and intrusion, thus identifying them as potential therapeutical targets to treat thought intrusion disorders.

Finally, our decoding analysis revealed that the content of the successfully suppressed thoughts is stored in the left IFC and the right LOC. Targeting these areas, with non-invasive brain stimulation (Sparing 2008) could help to minimize the unwanted effects of suppressed thoughts even when, subjectively, suppressed thoughts are successfully censored.

We believe that these results will shed light on the mechanisms of involuntary visual thoughts and the pervasiveness of their effects despite subjectively successful suppression by informing therapeutical strategies to prevent unwanted visual thought intrusions.

## Materials and Methods

### Participants

Experimental procedures were approved by the University of New South Wales Human Research Ethics Committee (HREC#: HC12030). All methods in this study were performed in accordance with the guidelines and regulations from the Australian National Statement on Ethical Conduct in Human Research (https://www.nhmrc.gov.au/guidelines-publications/e72). All participants gave informed written consent to participate in the experiment. We tested 15 participants, 4 women, aged 29.6±1.4 years old (mean±SEM). For analyses discriminating successful from failed suppression trials, a subset of 8 participant (2 women, aged 32±2.1 years old) was considered using as selection criterion of having at least 25% of failed or successful suppression trials.

We selected the sample size based on previous studies (Koenig-Robert and Pearson 2019; Soon et al. 2008; Aso et al. 2016) to meet standard criteria of statistical power. We conducted a post-hoc power analyses to ascertain the power achieved using GPower (Erdfelder et al. 2009). For GLM analyses, the power achieved was at least 0.91 at the voxel level (N=15, Figures 2 and 3). For the decoding analysis discriminating the content of successful suppression (Figure 4), we achieved a power of at least 0.79 to detect differences in the paired t-test for relevant conditions (N=8).

### Functional and structural MRI parameters

Scans were performed at the Neuroscience Research Australia (NeuRA) facility, Sydney, Australia, in a Philips 3T Achieva TX MRI scanner using a 32-channel head coil. Structural images were acquired using turbo field echo (TFE) sequence consisting in 256 T1-weighted sagittal slices covering the whole brain (flip angle 8 deg, matrix size = 256×256, voxel size = 1mm isotropic). Functional T2*-weighted images were acquired using echo planar imaging (EPI) sequence with 31 slices (flip angle = 90 deg, matrix size = 240×240, voxel size = 3mm isotropic, TR = 2000ms, TE = 40ms).

### Suppression/imagery task

We adapted the behavioural task from a previous study from our group (Kwok et al. 2018) to satisfy fMRI requirements. We instructed participants to either imagine or suppress (avoid imagining) the visual thought of either a red apple or a green broccoli (see Figure 1A). Each trial started with a written object cue reading “green broccoli” or “red apple” for 2s. After this, a task cue was shown, reading either “imagine” or “suppress” for 2s. Participants were instructed to either visualize as vividly as they could the cued object in the imagine condition or avoid thinking about the cued object in the suppress condition for 12s during which. A fixation point was shown on the screen and participants were required to fixate. In the suppression condition, participants were instructed to press a button as soon as they subjectively detected that the visual thought of the to-be suppressed object appeared in their minds. We labelled such events as “suppression breaks” and the trial was labelled as a failed suppression trail. The suppression break button could be pressed multiple times within the 12s (Supplementary Figure S4), thus representing multiple instantiations of suppression breaks. After the imagery/suppression period, a prompt asking to rate vividness (from 1 to 4, with 4 representing strongest vividness) was presented after imagery or failed suppressed trials. Participants responded by pressing one of the 4 buttons on two response boxes. No vividness question was shown after successful suppression trials which were automatically labelled as having vividness=0. In failed suppressed trials, whenever multiple suppression breaks were reported, we instructed participants to rate the highest vividness suppression break event. After reporting the vividness of the visual thought (if required), an inter trial interval of 10s was observed during which the word “rest” appeared on the screen. In each run of 5 min, 3 trials of each type (imagine/suppress apple/broccoli) were tested yielding a total of 12 trials. Trials were pseudorandomized within a run.

### Perception task

We presented flickering natural images of a broccoli and an apple against a black background at 4.167 Hz with 3 different perceptual intensities (40, 60 and 80% transparency) in order to maximize subsequent classifier generalization ability (Bannert and Bartels 2013). Natural images of a broccoli and an apple against a black background were retrieved on Google images search for images labelled for reuse with modification, and were presented inside a rectangle (the same that was used in the imagery/suppression task, Figure 1) including a fixation point at the centre. Within a run of 3 minutes, we presented the flickering imagers in a block manner, interleaved with fixation periods of 15 seconds each (i.e., apple: 15s, rest: 15s, broccoli: 15s, rest: 15s, etc.) Importantly, an attention task was performed consisting of detecting a change in fixation point brightness (+70% for 200ms). Fixation changes were allocated randomly during a run, from 1 to 4 instances. Participants were instructed to press any of the 4 buttons as soon as they detected the changes. Participants showed high performance in the detection task (d-prime=2.89 ±0.15 SEM).

### Functional mapping of retinotopic visual areas

To functionally determine the boundaries of visual areas from V1 to V4 independently for each participant, we used the phase-encoding method (Sereno et al. 1995; Warnking et al. 2002). Double wedges containing dynamic coloured patterns cycled through 10 rotations in 10min (retinotopic stimulation frequency = 0.033 Hz). To ensure deployment of attention to the stimulus during the mapping, participants performed a detection task: pressing a button upon seeing a grey dot anywhere on the wedges.

### Experimental procedures

We performed the 3 experiments in a single scanning session lasting about 1.5h. Stimuli were delivered using an 18” MRI-compatible LCD screen (Philips ERD-2, 60Hz refresh rate) located at the end of the bore. Participants held one dual-button response box in each hand (Lumina, Cedrus, San Pedro, CA, USA) that was used to record all responses. All stimuli were delivered and responses gathered employing the Psychtoolbox 3 (Brainard 1997; Pelli 1997) for MATLAB (The MathWorks Inc., Natick, MA, USA) using in-house scripts. Participants’ heads were restrained using foam pads and adhesive tape. Each session followed the same structure: first the structural scanning followed by the retinotopic mapping (10min). Then, the perception task was alternated with the imagery/suppression task until completing 3 runs of the perception task (3 min per run). Then the imagery/suppression task was repeated until completing 8 runs in total (5 min per run). Pauses were assigned in between the runs. The 4 first volumes of each functional runs were discarded to account for the equilibrium magnetization time and each functional run started with 10 seconds of fixation.

### Visual ROI functional definition

Functional MRI retinotopic mapping data were analysed using the Fast-Fourier Transform (FFT) in MATLAB. The FFT was applied voxel-wise across time points. The complex output of the FFT contained both the amplitude and phase information of sinusoidal components of the BOLD signal. Phase information at the frequency of stimulation (0.033Hz) was then extracted, using its amplitude as threshold (≥2 SNR) and overlaid them on each participant’s cortical surface reconstruction obtained using Freesurfer (Fischl, Sereno, and Dale 1999; Fischl et al. 2004). We manually delineated boundaries between retinotopic areas on the flattened surface around the occipital pole by identifying voxels showing phase reversals in the polar angle map, representing the horizontal and vertical visual meridians. In all participants, we clearly defined four distinct visual areas: V1, V2, V3, and V4. All four retinotopic labels were then defined as the intersection with the perceptual blocks (grating>fixation, p<0.001, FDR corrected) thus restricting the ROI to the foveal representation of each visual area.

### Suppressed-object information containing ROI definition

Areas bearing information about the content (apple vs broccoli) of the suppressed trials were defined using a decoding approach. Regressors for apple and broccoli were extracted from every run from the suppression trials (independent of success) using 12s boxcars. We used a leave one-run out cross-validation scheme (see MVPA subsection for details) and a searchlight approach. ROI containing information about the contents of suppression were defined as those a reaching a significant classification of Z > 2 (one-sample t-test against chance: 50%) at the voxel level. We then corrected for multiple comparisons using cluster-extent based thresholding employing Gaussian Random Field theory at p < 0.05. Two ROI were thus defined by this analysis, one in the inferior frontal gyrus and one in the lateral occipital cortex (Figure 4). Importantly, this ROI definition is orthogonal to the target decoding analysis where we tested the mutual informational content between perception and successful suppression as well as between imagery and suppression, as the training sets (perception and imagery trials) are independent from the training set (suppression trials) used to define the ROI.

### Lateralization analysis

We employed a classic measure of lateralization index (Oldfield 1971; Adcock et al. 2003). Hemispheric specific activations were extracted from spatially MNI normalized SPM beta volumes using right and left hemisphere masks. For every participant, a lateralization index (*LI*) was calculated as follows:

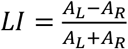; Where *A*_*L*_ and *A*_*R*_ are the sum of the activated voxels in the left and right hemisphere, respectively.

Thus, *LI* = −1 represents fully *left*-lateralized effects, while *LI* = +1, fully *right*-lateralized effects.

### Functional MRI signal pre-processing

All data were analysed using SPM12 (Statistical Parametric Mapping; Wellcome Trust Centre for Neuroimaging, London, UK). We realigned functional images to the first functional volume and high-pass filtered (128 seconds) to remove low-frequency drifts in the signal.

### Imagery vs suppression GLM analysis

Data were spatially normalized into the Montreal Neurological Institute (MNI) template and spatially smoothed using a FWHM 8mm 3D Gaussian kernel. We generated regressors for each condition (imagery and suppression, independent of the imagined/suppressed object) for each run independently. We used boxcar functions of 12s to model each trial with the canonical HRF as basis function. Vividness of the imagery and suppression trials was modelled using parametric modulators. General linear models (GLM) were used to test differences between imagery and suppression conditions. Participants’ (N=15) estimates (betas) of the mass-univariate GLM were fed into a second-level two-sample t-test analysis.

### Successful vs failed suppression GLM analysis

The analysis was performed as described in the previous paragraph except for the following differences. We generated regressors for successful and failed suppression (independent of the suppressed object) for each run independently. We used boxcar functions of 12s and 1s to model successful suppression and suppression breaks respectively, in order to capture their respective sustained and transient natures (Mitchell et al. 2007). Only participants having at least 25% of successful or failed suppression trials (N=8) were considered in order to having enough data to estimate the parameters.

### GLM analysis for multi-voxel pattern analysis

Data were analysed in their native space, without spatial normalization nor smoothing in order to avoiding disrupting information contained in the spatial patterns of activation (Hebart, Görgen, and Haynes 2014). For the task decoding (Figures 2C and 3B), we estimated GLM for imagery vs suppression and successful vs failed suppression as described above. For the content decoding (Figure 4), regressors for apple and broccoli were estimated using boxcar functions (15s for the perception trials 12s for the imagery and successful suppression conditions). This analysis was performed on the subset of participants having at least 25% of successful or failed suppression trials (N=8).

### Multi-voxel pattern analysis (MVPA)

We used a well-established decoding approach to extract information related to each grating contained in the pattern of activation across voxels of a given participant using the decoding toolbox (TDT) (Hebart, Görgen, and Haynes 2014). For the task decoding (Figures 2C and 3B), we used a leave one-run out cross-validation scheme. We trained a linear supporting vector machine (SVM) on all runs but one and then tested on the remaining one. We repeated this procedure until all runs were used as test and then averaged the results across validations (8-fold). Using this approach we tested whether information about the task nature could be decoded from functionally defined visual areas (from V1 to V4, see “Visual ROI functional definition” subsection for details). For the content decoding (Figure 4), we employed cross-classification to generalize information between perception or imagery and the successful suppression trials. We thus trained on the ensemble of the perception or imagery runs and tested on the ensemble of the imagery runs. No cross-validation was used here as the datasets were independent thus there was no risk of overfitting. We employed a region-of-interest (ROI) to test common representational content in functionally defined areas as containing suppressed object information (see “Suppressed-object information containing ROI definition” subsection for details). Decoding accuracies were averaged across runs and tested against chance level (50%) using a one-sample t-test across subjects.

### Statistical analysis on brain images

All second-level (across subjects) brain statistical images (derived from SPM or decoding) were subjected to a threshold at the voxel level p<0.001, as recommended in previous studies (Woo, Krishnan, and Wager 2014). We then corrected for multiple comparisons using cluster-extent based thresholding employing Gaussian Random Field theory (Friston et al. 1994; Worsley et al. 1996) at p<0.05, as implemented in FSL (Smith et al. 2004). Importantly, these thresholds have been shown to be valid within the nominal false positive ratios (Eklund, Nichols, and Knutsson 2016).

## Supplementary Figures

**Supplementary Figure S1.**
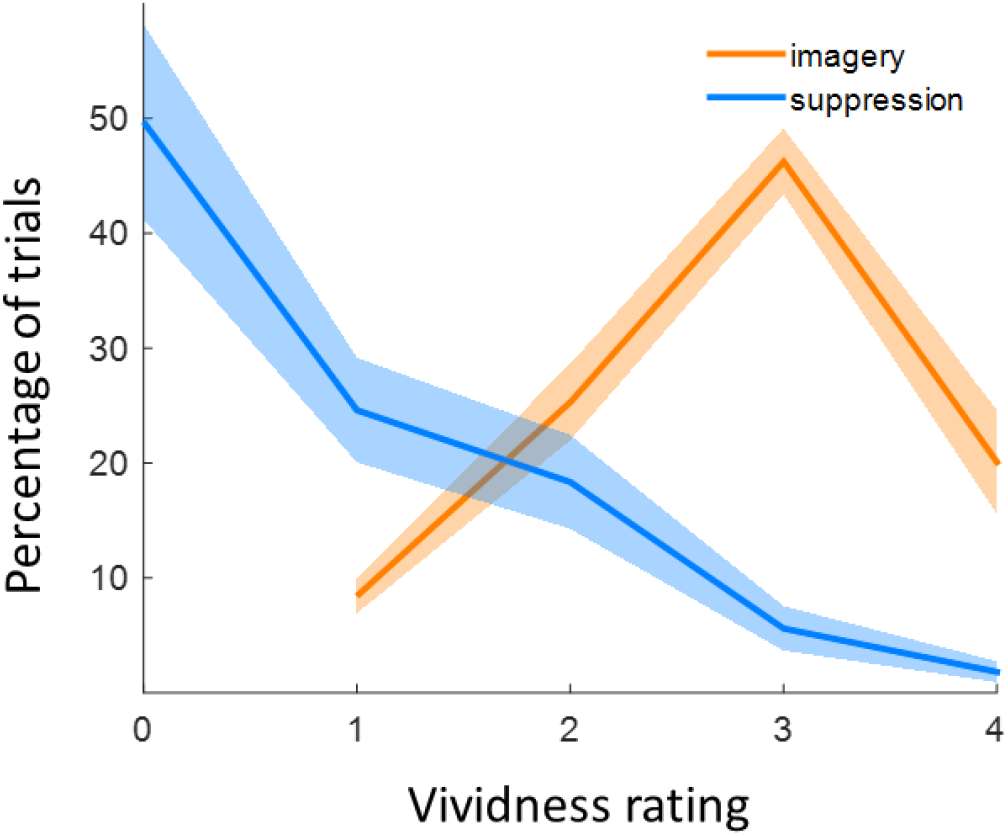
Vividness-rating distribution per condition. Subjective vividness ratings from 0 to 4. Zero represents successful suppression during the suppression condition. Orange line represents imagery while the blue represents suppression. Shaded area corresponds to the SEM across participants. Shade corresponds to ±1 SEM.

**Supplementary Figure S2.**
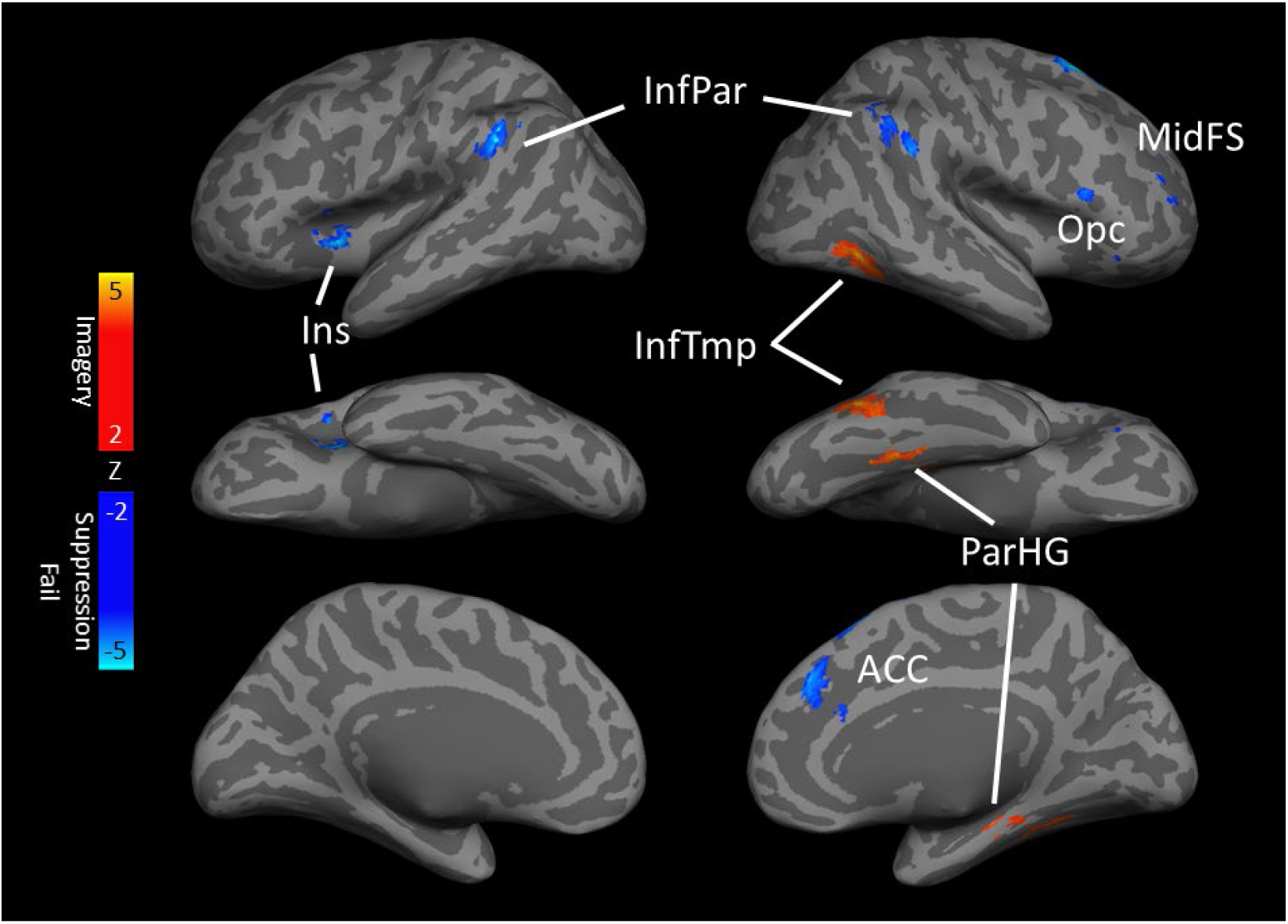
Imagery vs Suppression fail GLM. Comparing imagery and failed suppression revealed a similar network of areas as the contrast imagery vs suppression, except for a loss of lateralization of the suppressed activations compared to the previous contrast. All blobs are p<0.001 (voxel level) and p<0.05 cluster level correction (GRFT) for multiple comparisons. ACC: anterior cingulate cortex; InfPar: inferior parietal; InfTmp: inferior temporal; Ins: insula; MidFS: MidFS: middle frontal sulcus; ParHG: parahippocampal gyrus;

**Supplementary Figure S3.**
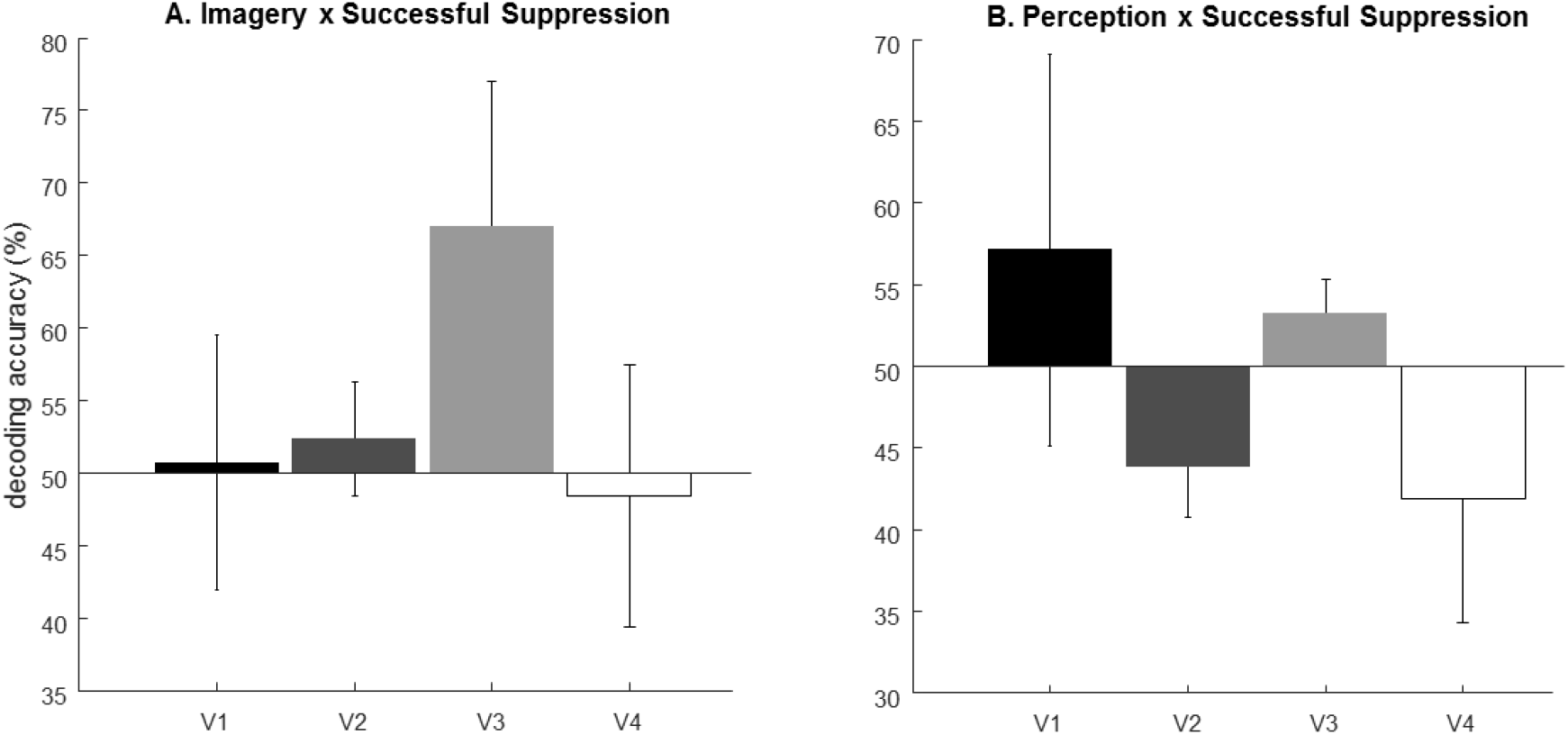
Visual areas cross-decoding. The content of successfully suppressed thoughts was not significantly decoded (p>0.05 one-tailed t-test) using information from imagery trials (panel A) nor from perception trials (panel B). Error bars correspond to ±1 SEM.

**Supplementary Figure S4.**
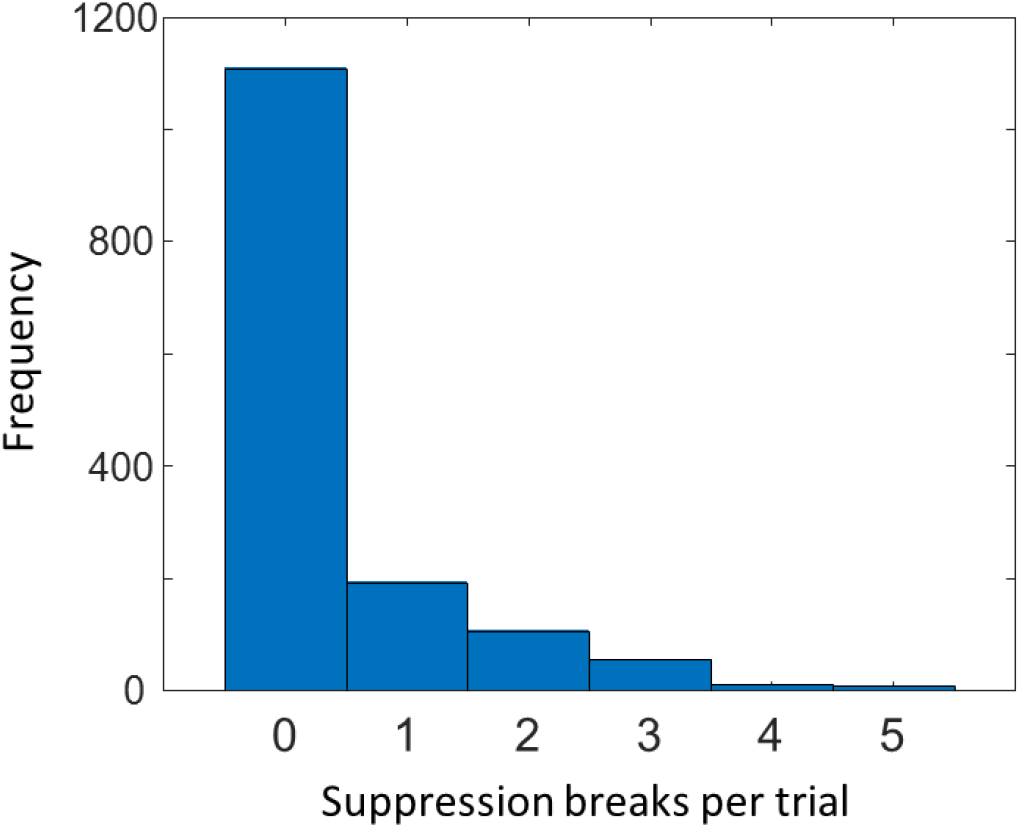
Histogram of suppression breaks per trial. Non-suppression breaks trials (0) outnumbered suppression break trials (1 to 5) by about 3:1.

